# In Vivo Structural Connectivity of the Reward System Along the Hippocampal Long-Axis

**DOI:** 10.1101/2023.09.11.556537

**Authors:** Blake L. Elliott, Raana Mohyee, Ian C. Ballard, Ingrid R. Olson, Lauren M. Ellman, Vishnu P. Murty

## Abstract

Recent work has identified a critical role for the hippocampus in reward-sensitive behaviors, including motivated memory, reinforcement learning, and decision-making. Animal histology and human functional neuroimaging have shown that brain regions involved in reward processing and motivation are more interconnected with the ventral/anterior hippocampus. However, direct evidence examining gradients of structural connectivity between reward regions and the hippocampus in humans is lacking. The present study used diffusion MRI and probabilistic tractography to quantify the structural connectivity of the hippocampus with key reward processing regions in vivo. Using a large sample of subjects (N=628) from the human connectome dMRI data release, we found that connectivity profiles with the hippocampus varied widely between different regions of the reward circuit. While the dopaminergic midbrain (VTA) showed stronger connectivity with the anterior versus posterior hippocampus, the vmPFC showed stronger connectivity with the posterior hippocampus. The nucleus accumbens (NAc) showed a more homogeneous connectivity profile along the hippocampal long-axis. This is the first study to generate a probabilistic atlas of the hippocampal structural connectivity with reward-related networks, which is essential to investigating how these circuits contribute to normative adaptive behavior and maladaptive behaviors in psychiatric illness. These findings describe nuanced structural connectivity that sets the foundation to better understand how the hippocampus influences reward-guided behavior in humans.

## Introduction

The hippocampus is a deep brain structure associated with episodic memory and spatial navigation (Burgess et al., 2002; Scoville & Milner, 1957; Squire, 1992; Lisman et al., 2017; Moser et al., 2008; O’Keefe & Nadel, 1978). However, the hippocampus is also functionally involved in a variety of adaptive behaviors including future planning (Buckner, 2010; Mullaly & Maguire, 2014; Stachenfeld et al., 2017; Benoit et al., 2011), reward-learning (Wirth et al., 2009; Devenport et al., 1981; Holscher et al., 2003; Ploghaus, et al. 2000; Ballard et al., 2019), novelty coding (Whittman et al., 2007; Bunzeck & Düzel, 2006; Kumaran & Maguire, 2007) and value-based decision-making (Palombo et al., 2015; Tang et al., 2021; Shadlen & Shohamy, 2016; Shohamy & Daw, 2015; Bunzeck et al., 2010; Wimmer & Shohamy, 2012). Its role in adaptive behavior has been mirrored in studies showing hippocampal alterations in maladaptive behaviors associated with reward processing, such as anhedonia and depression (Lodge & Grace, 2008; 2011; Grace, 2016; Lee et al., 2012; Zackova et al., 2021). While the focus of this prior work has been centered on the hippocampus, evidence suggests it subserves these adaptive behaviors through structural and functional connectivity with brain regions guiding motivation. Non-human animal histology has shown that brain regions involved in reward processing and motivation, including the ventromedial prefrontal cortex (vmPFC), nucleus accumbens (NAc), and ventral tegmental area (VTA), are interconnected with the hippocampus (Lisman & Grace, 2005; Haber & Knutson, 2010; Poppenk et al., 2013). However, evidence for this structural connectivity in humans is lacking, leaving open questions about whether these circuits show profiles homologous to those seen in rodents and non-human primates. Here, we characterized structural connectivity of reward-related regions across the long-axis of the hippocampus in humans using diffusion MRI.

Motivation is a crucial factor driving learning and goal-directed behavior. Motivation is mediated by engaging a network of brain regions (Haber & Knutson, 2010), centered on the ventral tegmental area (VTA), nucleus accumbens (NAc), and medial prefrontal cortex (mPFC), particularly its more ventral portions (e.g., ventromedial prefrontal cortex, vmPFC), suggesting that they are prime targets to investigate in the context of hippocampal structural connectivity. The VTA consist of dopamine neurons that send reward prediction error signals (along with other heterogenous signals, Matsumoto & Hikosaka, 2009; Bromberg-Martin et al., 2010; Lammel et al., 2014; Ljungberg et al., 1992) to the striatum and other brain regions. The NAc is critical for associative learning and motivation (Mogenson et al., 1980; Yang et al, 2018; Schultz et al., 1997). The vmPFC supports reward learning and decision-making by representing abstracted reward or affective value and providing predictions about future outcomes (Rolls, 2019; Rudebeck & Murray 2014; Wilson et al. 2014). Critically, research in animals has shown the hippocampus to have direct and/or indirect connectivity to each of these regions (Kelley et al., 1982; Groenewegen et al., 1987; Scofield et al., 2016; Cavada et al., 2000; Fanselow & Dong, 2010; Gasbarri et al., 1991; Gasbarri et al., 1996; Swanson et al., 1982).

Connectivity with the hippocampus is not homogenous, but rather the dorsal and ventral hippocampus –which corresponds with the posterior and anterior hippocampus in humans, respectively– have distinct patterns of connectivity with other brain regions, which may underlie their functional differences. Specifically, the dorsal/posterior hippocampus is more strongly connected with parietal and retrosplenial cortices, reflecting its role in navigation, reinstatement, and retrieval (Moser & Moser, 1998; Sherril et al., 2013; Whitlock et al., 2008; Kim, 2015; Sheldon & Levine, 2016). The ventral/anterior hippocampus is more connected with the amygdala, striatum, hypothalamus, and medial prefrontal regions (Fanselow & Dong, 2010; Strange et al., 2014), reflecting its role in affective processing, emotion regulation, and emotional memory (LeDoux, 1993; Mather, 2007; Phelps, 2004; Zhu et al., 2019). These structural differences have functional consequences, as evidenced by studies showing that dorsal hippocampal structure is correlated with spatial memory performance, whereas ventral hippocampal structure is correlated with emotion regulation (Bannerman et al., 2003; Bannerman et al., 2004; Nadel, 1968; Woollett & Maguire, 2011; Van Rooij et al., 2015; Willard et al., 2009; Richardson et al., 2004; Snytte et al., 2022; Moser & Moser, 1998). These patterns of connectivity suggest that the flexibility of different adaptive behaviors may result in differential connectivity of reward regions along the hippocampal long axis, however, how reward regions preferentially target the hippocampus has yet to be dissected in humans.

Although tracer studies and functional imaging in humans have begun to classify the organization of the HPC long-axis, in-vivo structural connectivity in humans is lacking. Here, we leveraged a large diffusion weighted imaging (DWI) dataset collected as part of the Human Connectome Project and utilized probabilistic tractography to segment the hippocampus according to its connectivity to regions associated with motivation and adaptive behavior, including the VTA, NAc, and vmPFC. Further, we generated a probabilistic atlas of the hippocampus reflecting how connectivity with reward regions vary across its long-axis to bolster research on hippocampal contributions to adaptive behavior. Delineating the structural connectivity of the hippocampal long-axis with reward regions in humans is crucial for interpreting human neuroimaging findings and translating findings from animal research, as well as understanding the role of these regions in adaptive and maladaptive behaviors.

## Method

### Participants

The study sample comprised participants enrolled in the Human Connectome Project (HCP). Data were obtained from the WU-Minn HPC Consortium S900 Release; participants from whom T1-weighted and diffusion-weighted MRI scans were acquired, and for whom complete structural metrics generated using the HCP FreeSurfer pipeline, were included in this study (WU-Minn HCP Consortium, 2015). Altogether, 628 participants were included in the final analyses.

### MRI Acquisition, Preprocessing, and Analysis

HCP data acquisition and preprocessing pipeline procedures are detailed here: (Van Essen et al., 2013, Van Essen et al., 2012, Barch et al., 2013, Glasser et al., 2013). We utilized the minimally preprocessed diffusion MRI (dMRI) data that were provided by the HCP S900 release. The dMRI data had gone through EPI distortion, eddy current, and motion correction, gradient nonlinearity correction, and registration of the mean b0 volume to a native T1 volume. In addition to the HCP minimally pre-processed pipeline, we processed the dMRI data with FSL’s BEDPOSTX (Behrens et al., 2007) to model white matter fiber orientations and crossing fibers.

### Structural connectivity analysis

#### Tractography

The diffusion-weighted data were processed using FSL FDT toolbox (www.fmrib.ox.ac.uk/fsl). Measures of tract strength were calculated using probabilistic tractography with a partial volume model (Behrens et al., 2003), allowing for up to 2 fiber directions in each voxel (Behrens et al., 2007). Fiber tracking was conducted in parallel for each voxel within a predefined seed mask (bilateral hippocampus). We used 5,000 samples per voxel, a curvature threshold of 0.2, and a step length of 0.5 mm. Tractography analyses were conducted in each subject’s native anatomical space and the results registered to Montreal Neurological Institute (MNI) space by providing transformation parameters estimated via a 2-step procedure. First, the fractional anisotropy (FA image) was registered to each subject’s high-resolution T1- weighted image using FSL’s linear image registration tool with six degrees of freedom and a mutual information cost function (FLIRT; Jenkinson et al., 2002). Next, the T1-weighted image was nonlinearly registered to the 2 × 2 × 2 mm^3^ nonlinear MNI template with FSL’s non-linear image registration tool (FNIRT). Finally, the transformation parameters obtained from these two steps were concatenated to yield the mapping from the DWI to MNI space. Tractography was performed separately for the left and right hippocampus, and possible tracts were restricted to the hemisphere of origin using an exclusion mask of the contralateral hemisphere.

#### Segmentation

To assess connectivity of the hippocampus with reward-related regions, we used four masks, defined *a priori*. Following previous diffusion tractography segmentations of the thalamus (Behrens et al., 2003; Johansen-Berg et al., 2005) and striatum (Tziortzi et al., 2014) the target regions of interest were chosen based on based on their anatomical characteristics. It’s important to note that these anatomical regions naturally differ in size. This size variation could potentially influence general observations, but it’s more relevant for analyzing individual differences in tract-strength rather than creating distinct within-subject parcellations. Our primary focus centers on exploring interactions along the longitudinal axis of the hippocampus. In this context, any potential effects related to the size of these regions of interest are unrelated.

The seed masks (hippocampus) were defined using the Harvard-Oxford subcortical atlas integrated within FSL (Figure 1). The target area of the limbic (ventral) striatum was defined using a connectivity-based segmentation atlas with subdivisions for sensorimotor, executive, and limbic regions (Tziortzi et al., 2014). The vmPFC masks were defined from Bhanji et al. (2019). This mask is an inclusive vmPFC/OFC mask that includes the whole area of prefrontal cortex that is both ventral (z < 0 in standardized coordinate space) and medial (i.e., superior and inferior medial gyri, anterior cingulate gyrus, gyrus rectus, medial orbital gyrus, and the adjacent sulci). The functional relevance of this mask was investigated using the Neurosynth meta-analytic engine and topic-based mapping (Yarkoni et al., 2011; Poldrack et al., 2012). Bhanji et al. discovered that vmPFC activation was significantly associated with social, emotion, and decision-making functions (2019). The seed ROI for the SN/VTA was defined using a probabilistic atlas of human SN/VTA (Murty et al., 2014). We used a 50% probability threshold. The MNI space target masks were normalized to each participant’s native space using the inverse of the spatial normalization parameters. To tailor the ROIs to individual anatomy, we masked the ROIs with individually segmented gray matter (GM) images generated from freesurfer.

**Figure 1:**
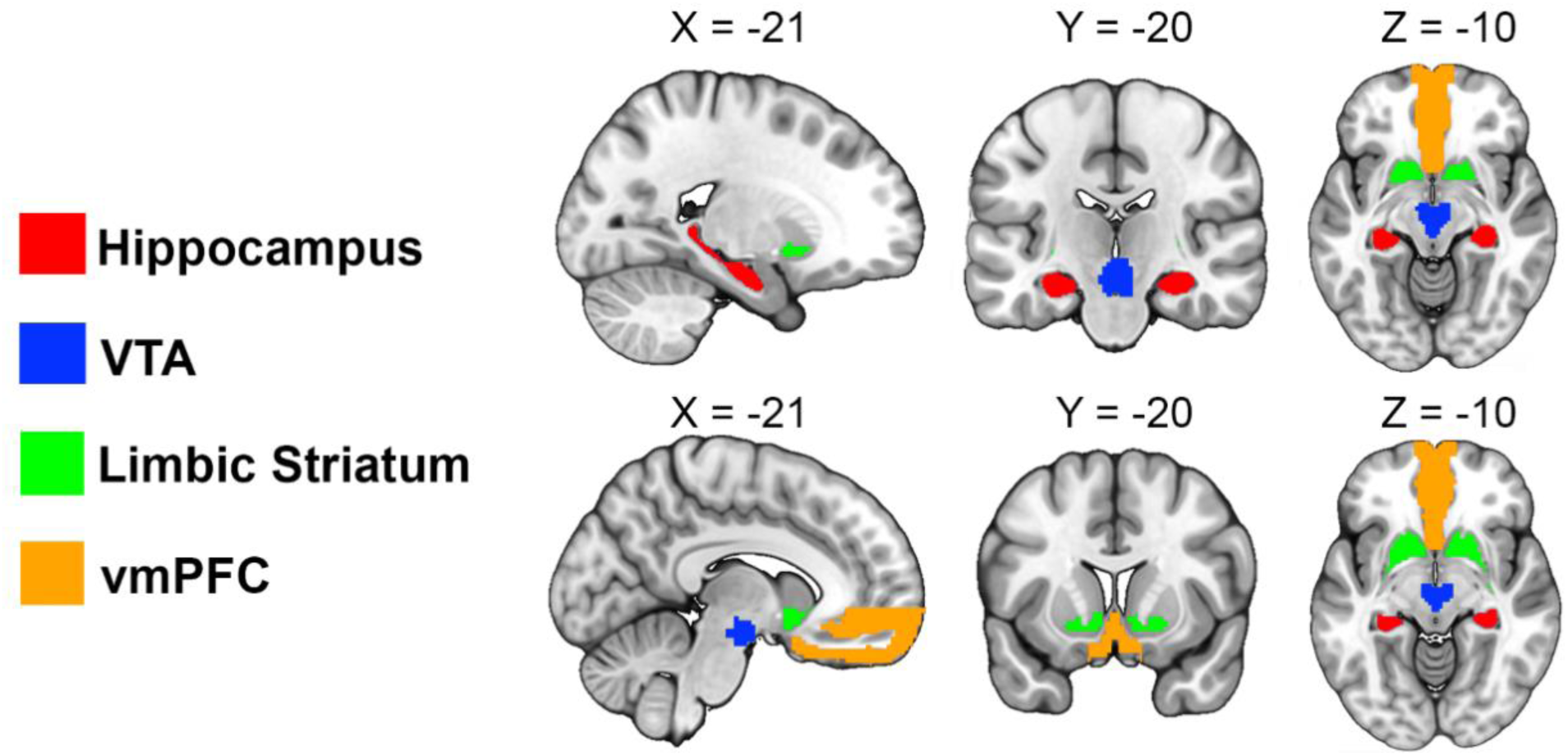
Seed and target regions for the probabilistic tractography analysis. The target region of the limbic striatum was functionally segmented based on projections from motor, executive, and limbic cortices (Tziortzi et al., 2014). The target region of the vmPFC was functionally correlated with meta-analyses of social, emotion, and economic valuation or decision-making (Bhanji et al., 2016). The target region of the midbrain was generated from VTA probabilistic atlas (Murty et al., 2014). The seed region of the hippocampus was generated from the Harvard-Oxford subcortical atlas (Frazier et al., 2005).

**Figure 2:**
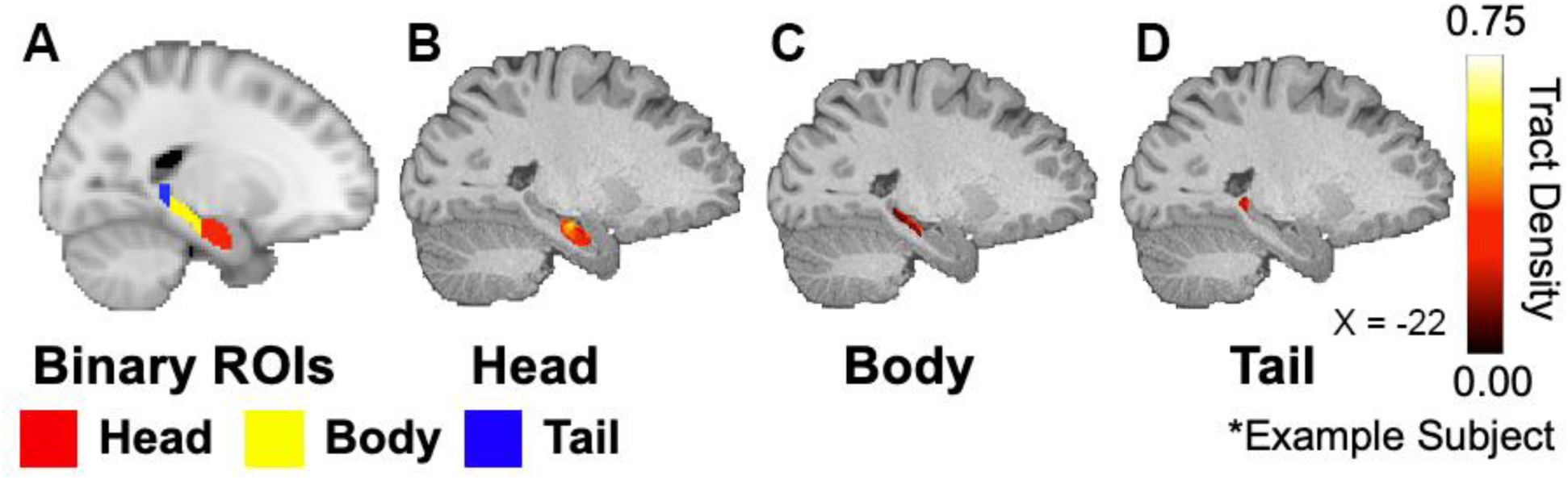
Within-subject tract density measure along the hippocampal long axis (head, body, and tail). A) Head, body and tail ROIs in MNI space. B) Voxel-wise tract density in the head of the hippocampus. C) Voxel-wise tract density in the body of the hippocampus. D) Voxel-wise tract density in the tail of the hippocampus. Mean voxel-wise tract density in each hippocampus ROI was computed for use in the ANOVA.

Projections from the hippocampus to the vmPFC, limbic striatum, and VTA were estimated following standard procedures, such that seed-based classification maps were first thresholded so that only voxels with at least 10 tracts terminating in one of the target regions were kept (Cohen et al., 2009; Forstmann et al., 2012, van den Bos, 2014). Next, the voxel values were converted into proportions of the number of tracts reaching the target mask from one voxel, divided by the number of tracts generated from that voxel (maximum 5,000).

#### Hippocampus Topology

To assess the topology of connectivity of reward regions along the hippocampal long-axis, each subject’s value map was divided into head, body, and tail regions (Duvernoy et al., 1998). Notably, the head represents more ventral/anterior portions whereas the tail represents more dorsal/posterior portions. We used the mean of these value maps as the measure hippocampal connectivity across its long axis with the SN/VTA, NAc, and OFC. Differences in the topology of hippocampus connectivity along the long-axis were using a repeated measures Analysis of Variance (ANOVA) with the following within-subject factors: reward region (vmPFC, limbic striatum, VTA), hippocampal long-axis region (head, body, tail) and hemisphere (left, right). We tested for differences between specific reward regions and long-axis regions using pairwise comparisons with Bonferroni correction. R (version 4.1.2) was used for all analyses.

The head, body, and tail of the hippocampus were defined in the MNI 152 T1-weighted image (Fonov et al., 2009; Fonov et al., 2011) using the anatomical benchmarks outlined by the Hippocampal Subfields Group for the European Joint Programme for Neurodegenerative Disease Research (JPND; Olsen et al., 2019). The hippocampal head comprises the region between (and including) the anterior most slice (in the coronal view) in which the hippocampus appears and the posterior most slice in which the uncus is visible. The hippocampal body consists of the region between the hippocampal head and the last slice in which the lamina quadrigemina (colliculi) are visible in the posterior brain stem. The hippocampal tail comprises the region between the posterior-most slice of the body and the posterior-most slice in which the hippocampal formation is visible.

#### Hippocampus Connectivity Atlas

To generate the probabilistic hippocampus connectivity atlas, we created group-averaged maps with three levels of thresholding. Following standard procedures (Tziortzi et al., 2014; Elliott et al., 2022a) we thresholded each individual’s ROIs to the vmPFC, VTA, and limbic striatum at 50 streamlines per voxel for each hemisphere. Once each individuals’ ROIs were thresholded, they were binarized and averaged together to create a group-averaged atlas. The resulting atlases (HPC-Limbic Striatum, HPC-vmPFC, HPC-VTA) are publicly available at https://github.com/blelliott23/HCP-Hippocampus-Reward-Diffusion-Segmentation.

The previous method allows for voxels to overlap between each ROI (as long as the voxel met the minimum number of streamlines to that ROI). An additional “hard” segmentation was conducted following standard procedures (Johansen-Berg et al., 2005; Tziortzi et al., 2014). This segmentation precludes individual voxels from overlapping within a single subject (Fig. 4). To generate this “hard” segmentation, ROIs for each individual were thresholded at 10 streamlines. Next, each voxel was calculated as a proportion of the total number of streamlines from that voxel to reach any target. Each voxel was then assigned to the target region that had the highest probability of connection. These ROIs were then binarized and averaged together to create a group-averaged atlas. The resulting atlas (HPC-Limbic Striatum, HPC-vmPFC, HPC-VTA) are publicly available at https://github.com/blelliott23/HCP-Hippocampus-Reward-Diffusion-Segmentation.

## Results

### Hippocampus topology

Hippocampus connectivity to each reward region is summarized in **Figure 3**. A 3×3×2 repeated measures ANOVA with Greenhouse-Geisser corrections was conducted to investigate whether hippocampal connectivity (tract density) varied among reward regions (vmPFC, limbic striatum, VTA), long-axis region (Head, Body, Tail), and hemisphere (Left, Right), as well as their interactions.

**Figure 3:**
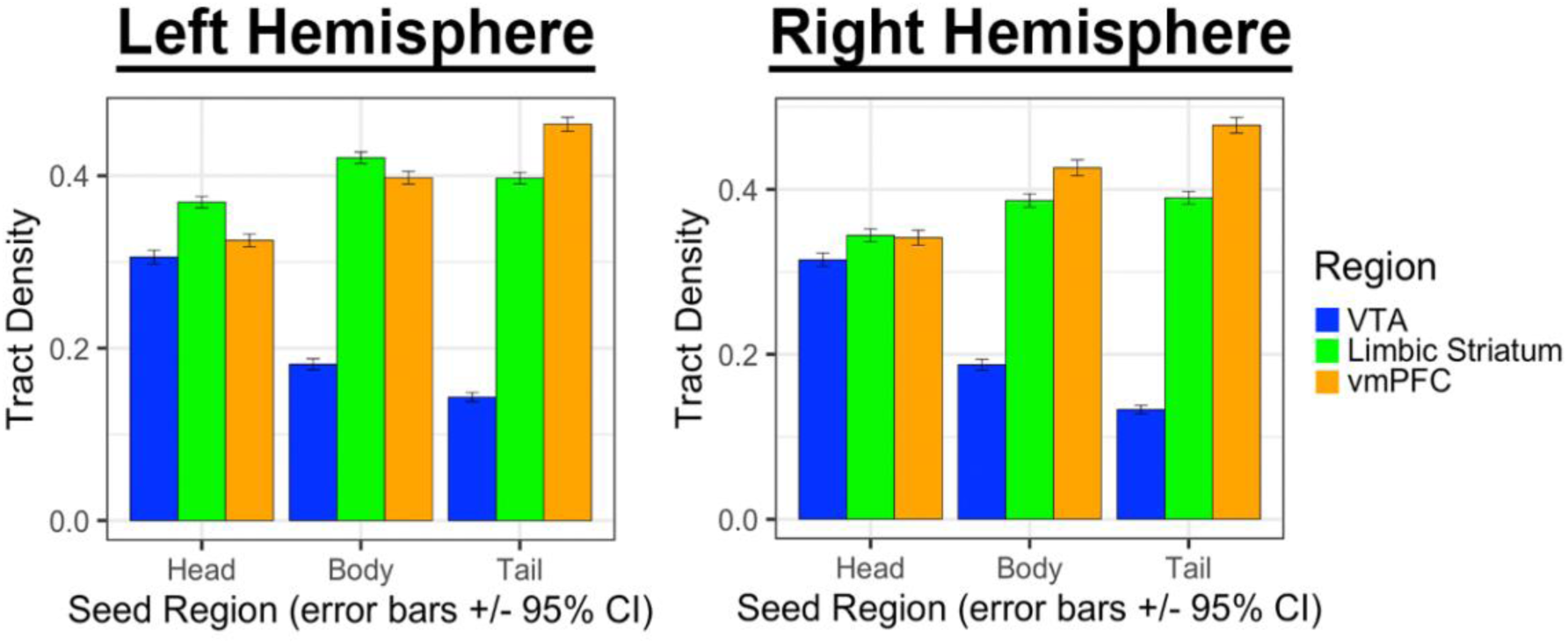
Within-subject mean tract density to each reward ROI along the hippocampal long axis (head, body, and tail) for each hemisphere. Error bars represent 95% confidence intervals. All main effects and interactions are statistically significant.

**Figure 4:**
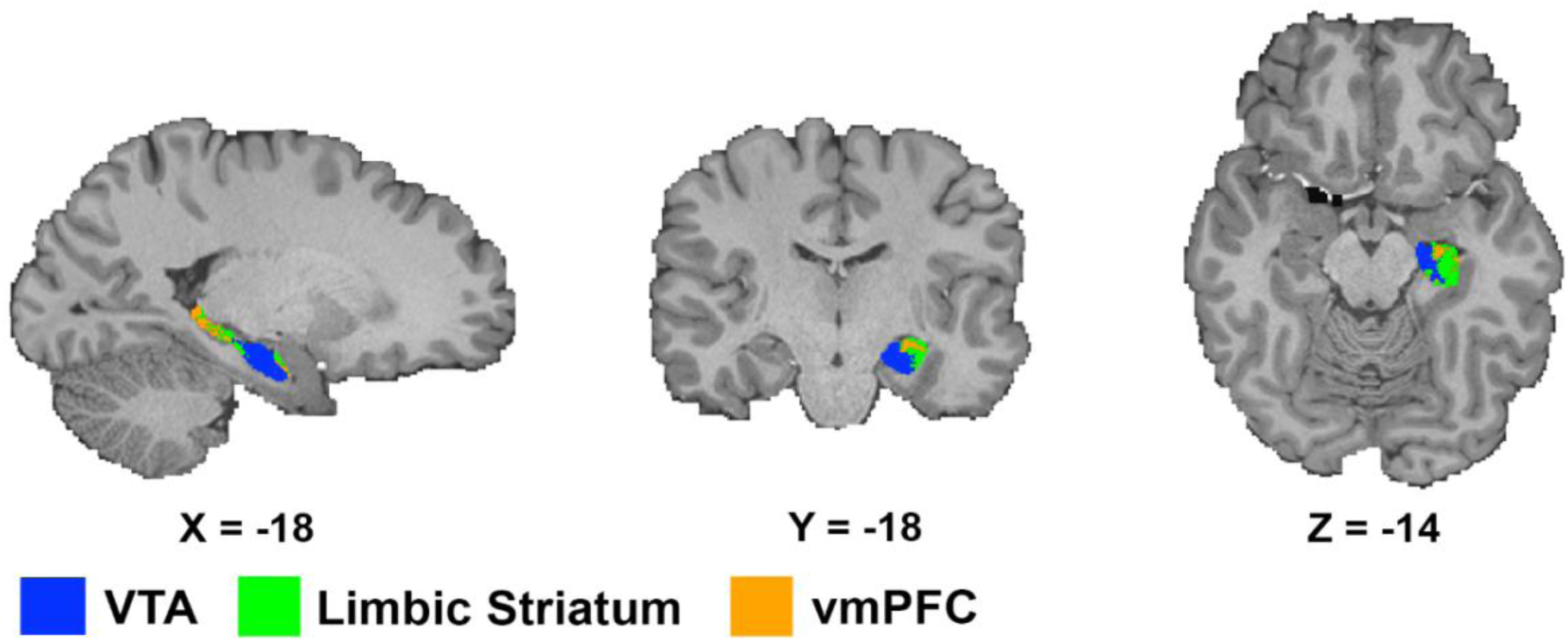
Hard segmentation on an example subject. The hippocampus was segmented by assigning each voxel to the lobe with which it had the highest connection probability (Johansen-Berg et al. 2005). After this “hard” segmentation, the areas in the hippocampus that associate with each target region were established.

### Main Effect of Reward region

There was a significant main effect of reward region, F(1.80, 1126.78) = 923.29, p < .001, ηp2 = .596, indicating that tract density differed significantly across the three reward regions (vmPFC, limbic striatum, and VTA). Post-hoc analyses with Bonferroni correction revealed that tract density for the vmPFC (M = 0.41, SD = 0.12) was significantly higher than the limbic striatum (M = 0.38, SD = 0.09, p < .001) and the VTA (M = 0.21, SD = 0.11, p < .001). Tract density for the limbic striatum was significantly lower than the vmPFC, but higher than the VTA (p < .001).

### Long-axis region

There was a significant main effect of hippocampal long-axis region (F(1.60, 1004.25) = 7.83, p < .001, ηp2 = .012) indicating that tract density differed significantly across the three long-axis regions (head, body, and tail). Post-hoc analyses with Bonferroni correction revealed that tract density for the head region (M = 0.33, SD = 0.10, p < .001) was significantly higher than the body (M = 0.33, SD = 0.14, p < .001, mean difference = 3.19e^-5^) but not significantly different from the tail (M = 0.33, SD = 0.17, N.S., mean difference = 5.76e^-7^). Tract density for the body was significantly lower than the head and the tail (mean difference = 3.25e^-5^ p = .001).

**Figure 5:**
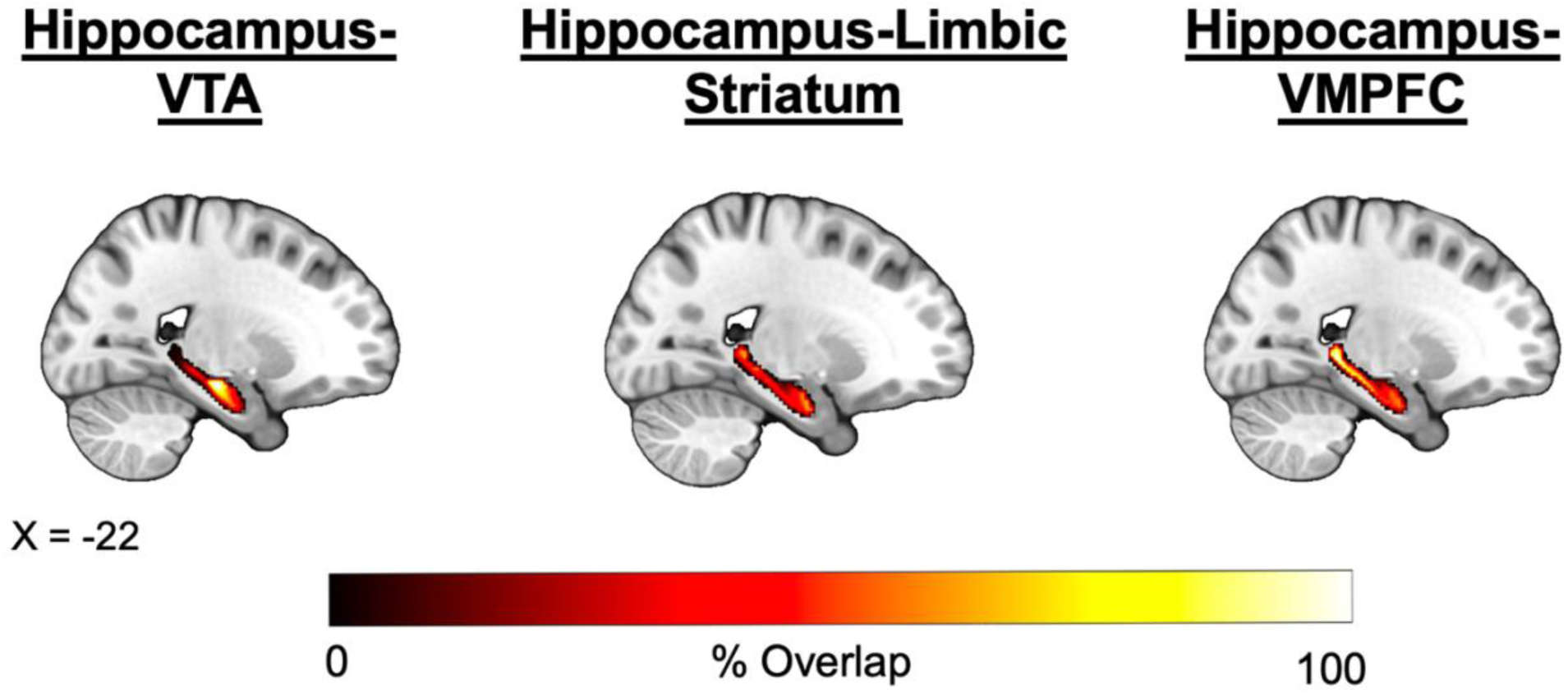
Hard segmentation group-averaged projections. Heat map represents the percent overlap of participants. A probabilistic atlas of the group-averaged projections is freely available on Github.

### Hemisphere

There was a small, but significant main effect of hemisphere, F(1, 627) = 19.02, p <. 001, ηp2 = .029. Tract density was relatively greater in the left compared to the right hemisphere (M = 0.33, SD = 0.14 vs. M = 0.33, SD = 0.13, p < .001, mean difference = 2.85e^-5^).

### Interactions

### Reward region x hippocampal long-axis region

There was a significant interaction between reward region (vmPFC, limbic striatum, VTA) and hippocampal long-axis region (head, body, tail), F(2.42, 1515.94) = 2475.38, p < .001, ηp2 = .798, indicating that tract density to each reward region differed across the three long-axis regions. Post-hoc analyses with Bonferroni correction revealed that the VTA connectivity with the head of the hippocampus (M = 0.31, SD = 0.10) was significantly higher than the body (M = 0.18, SD = 0.08, p < .001) and the tail region (M = 0.14, SD = 0.06). Tract density from the body was significantly higher than the tail (p < .001). The results suggest that in humans, the hippocampus has graded connectivity with the VTA, with the strongest connectivity from the anterior (head) region of the hippocampus.

Post-hoc analyses of hippocampal long-axis connectivity with the vmPFC from the head of the hippocampus (M = 0.33, SD = 0.10) was significantly lower than the body (M = 0.41, SD = 0.12, p < .001) and the tail region (M = 0.47, SD = 0.11). Tract density from the body was significantly lower than the tail (p < .001). The results suggest that in humans, the hippocampus has graded connectivity with the vmPFC, with the strongest connectivity from the posterior (tail) region of the hippocampus.

Post-hoc analyses of hippocampal long-axis connectivity with the limbic striatum revealed that the head of the hippocampus (M = 0.36, SD = 0.09) was significantly lower than the body (M = 0.40, SD = 0.09, p < .001) and the tail region (M = 0.39, SD = 0.01). Tract density from the body was significantly higher than the tail (p < .001). The results suggest that in humans, the hippocampus has a distributed connectivity profile with the limbic striatum, with a slight preference for the strongest connectivity from the body of the hippocampus.

## Discussion

This is the first study, to our knowledge, to delineate the structural connectivity of the hippocampal long-axis with key nodes across the reward circuit using diffusion MRI and probabilistic tractography in humans. We found that the hippocampus has a distinct connectivity profile across the long axis to each reward region examined. The dopaminergic midbrain (VTA) displayed the strongest connectivity to the anterior hippocampus. The vmPFC displayed stronger connectivity to the posterior hippocampus. Finally, the limbic striatum was more distributed along the hippocampal long-axis, with a slight preference for the body of the hippocampus. We compiled these connectivity profiles to generate a publicly available, probabilistic atlas of the hippocampus centered on structural connectivity with reward-related networks to support future neuroimaging studies characterizing hippocampal involvement in adaptive behaviors.

Although research in non-human animals has progressed in characterizing hippocampal-reward circuits, studies in humans are scarce, limiting our ability to translate findings from animal studies when considering hippocampal contributions to adaptive behavior. In this study, we probed hippocampal connectivity with three reward ROIs: The VTA, vmPFC, and nucleus accumbens (NAc). We selected these regions based on known non-human primate anatomy, and a human meta-analyses of functional neuroimaging studies using the terms "reward" and "subjective value" using two separate methods (an automated tool Neurosynth, Figure 6; and a researcher generated approach, Batra et al. 2013; Yarkoni et al., 2011; Poldrack et al., 2012). To our knowledge, our study is the first to characterize hippocampal long-axis connectivity to these regions. While we have provided the foundation for understanding the relationship between reward regions and the hippocampus here, it will be important in the future to extend these efforts to other nuclei and circuits that are involved in reward processing (Haber & Knutson, 2010).

**Figure 6:**
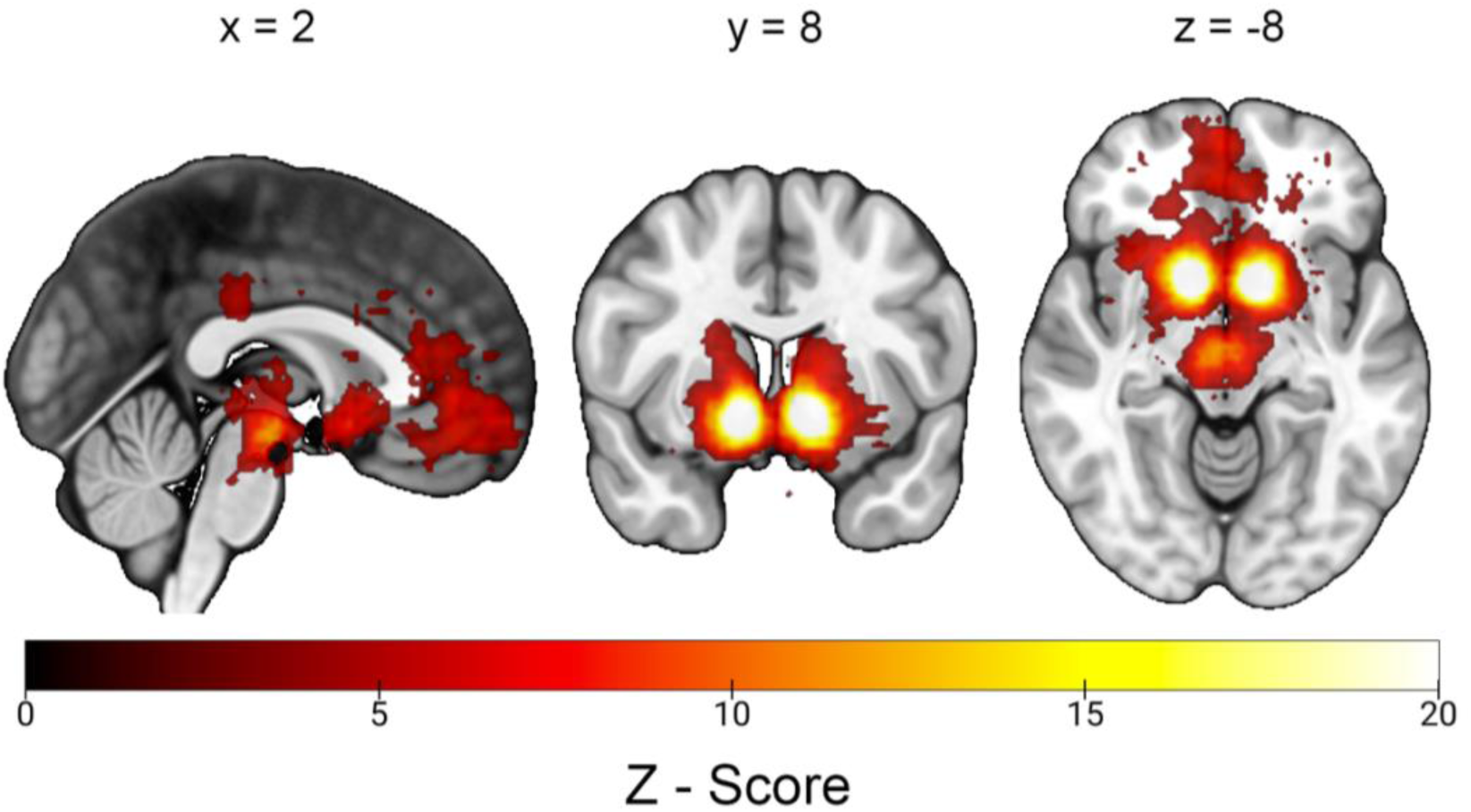
Neurosynth term-based meta-analysis conducted for the terms “reward”. The meta-analysis revealed large clusters centered on the VTA, ventral striatum, and vmPFC. The map can be examined at http://www.neurosynth.org/analyses/terms/reward.

The VTA forms a bi-directional circuit with the hippocampus which invigorates adaptive behavior in response to reward and novelty, as well as prioritizing memory for these events (Lisman et al., 2011; Rutishauser, 2019; Legault & Wise, 1999; Legault & Wise, 2001; Lodge & Grace, 2006), which is critical for behaviors such as reward memory and decision-making (Adcock et al, 2006, Elliott et al., 2020a; Elliott et al., 2020b; Shohamy & Adcock, 2010; Shohamy & Daw, 2015). Our results revealed that connectivity with the VTA was localized predominantly in the anterior hippocampus, which is consistent with rodent studies showing that VTA dopamine neurons predominantly innervate the ventral hippocampus (Gasbarri 1994a, 1994b; Oades & Halliday, 1987; Swanson, 1982; Verney et al., 1985).

These VTA findings results are also in line with human fMRI studies (Krebs et al., 2011; Adcock et al., 2006; Murty et al., 2017). Prior work using human neuroimaging demonstrated VTA-HPC connectivity to be critical for reward-motivated memory encoding (Adcock et al., 2006; Wittmann et al., 2005; Shigemune et al., 2014; Wolosin et al., 2012; Gruber et al., 2016), with studies showing a bias towards engagement of anterior hippocampus (Poppenk et al., 2013; Cowan et al., 2021). Further, previous research has found structural connectivity between the VTA and HPC to be positively correlated with individual differences in reward-motivated memory performance (Elliott et al., 2022b). Our findings integrate these two lines of research by providing a putative mechanism for functional biases towards the anterior hippocampus during motivated memory based on its structural connectivity, which supports future studies using multi-modal imaging approaches to understand function-structure relationships.

The HPC and vmPFC have been implicated in adaptive behaviors including reward-learning, motivation, decision-making, and episodic memory. In non-human primates HPC-vmPFC connectivity has been associated with prospection (Rolls 2019; Rudebeck & Murray 2014). The vmPFC represents abstracted reward or affective values and provides predictions about future outcomes (Rolls 2021; Rudebeck & Murray 2014; Mainen & Kepecs, 2009; Kable & Glimscher, 2009; Klein-Flugge et al., 2022). Previous research has shown that HPC-vmPFC connectivity is crucial for more adaptive forms of episodic memory, including remote autobiographical memory, schematic representations, and inference (Gilboa & Marlatte, 2017; McCormick et al., 2018; Schlichting & Preston, 2015). Notably, these functions have been hypothesized to rely primarily on anterior HPC-vmPFC activity (Abela & Chudasama, 2013; Schumacher et al, 2016; Viard 2011; Monk et al., 2021), which is consistent with structural connectivity we found between the most anterior portions of the hippocampus and vmPFC, but is surprising given the robust connectivity we found with the posterior portions of the hippocampus (discussed below). In fact, HPC-vmPFC connectivity from the head of HPC was greater than HPC-VTA connectivity from the head. This could be a potentially important area where dopaminergic reward signals from the VTA integrate with signals from the vmPFC to support adaptive behaviors. Additionally, visual inspection of the HPC-vmPFC atlases revealed that the most anterior portion of the HPC had greatest connectivity to the vmPFC and limbic striatum. This finding could be meaningful in terms of localizing function within the anterior HPC. However, we did show robust connectivity of the tail of the hippocampus (i.e., posterior hippocampus) with the HPC. This raises important questions about the role of the posterior hippocampus in adaptive behaviors and suggests that more attention should be paid to types of representations stored in the posterior hippocampus, both spatial and non-spatial, contribute to adaptive behavior.

Regarding the NAc, we found a relatively more homogenous connectivity profile across the hippocampal long axis. Animal work has shown that the hippocampus has strong efferent projections to the nucleus accumbens, via the ventral subiculum (Legault & Wise, 2001, Blaha et al., 1997; Taepavarapruk et al., 2000; Floresco et al., 2001; Floresco et al., 2003), which is known to regulate reward behavior (LeGates et al,. 2018), as well as stimulate reward seeking behavior in previously rewarded contexts. In line with these functions, rewards increase synchronization between HPC and NAc neurons (Tabuchi et al., 2000), which have been shown to be critical for drugs of abuse (Sjulson, et al., 2018). The NAc is also necessary to relay hippocampal signals to DA neurons (Lisman & Grace, 2005), which dovetails with human neuroimaging showing that HPC-NAc connectivity is associated with reward motivation and associative learning (Ballard et al., 2019; Shigemune et al., 2014) as well as interactions between feedback learning and episodic memory (Davidow et al., 2016). Additionally, resting state fMRI in humans has shown maximal correlation between the NAc and the body of the HPC (Kahn & Shohamy, 2013), consistent with our results. Given our structural connectivity results, we predict that NAc-hippocampal signaling may be implicated in a wide range of behaviors given that it has diffuse projections across the hippocampus.

Previous animal studies have found connectivity with the NAc from both the dorsal and ventral hippocampus (Groenewegen et al., 1996; Naber & Witter, 1998; Swanson & Kohler, 1986; Fanselow et al., 2010). However, it is thought that a gradient exists both structurally and functionally with the NAc along the hippocampal long-axis, such that more anterior regions innervate the medial NAc (shell) and are involved in more affective behaviors (Strange et al., 2014). In line with animal studies, our results found distributed and relatively homogenous NAc connectivity along the HPC long-axis. Although not investigated in the current study, it is possible that our results would differ should specific regions of the NAc (i.e. core and shell) be considered. Another exciting structure to examine is the fornix, which is known to provide connections to the NAc with the long-axis of the hippocampus. In line with our results, the fornix is distributed along the entire hippocampal long-axis, and has been shown to be intimately involved with both motivated behavior and the generation of hedonic responses (Trouche et al., 2019).

Prior work, outside the domain of reward, has shown that the hippocampus is anatomically and functionally distinct along its long-axis (ventral-dorsal in rodents, anterior-posterior in primates) both in its internal structure as well as its broader network connectivity. Within the hippocampus, the anterior and posterior hippocampus have distinct proportions of subfields (lower proportion of DG in anterior HPC than in posterior HPC, and a higher proportion of CA1–3 in anterior HPC than in posterior HPC, which may reflect differences in neurogenesis) as well as distinct cytoarchitecture and genetic domains (Fanselow & Dong, 2010; Dong et al., 2009; Thompson et al., 2008). Regarding network connectivity, in non-human primates, the posterior hippocampus has stronger connectivity with the retrosplenial cortex, area TE in the inferior temporal lobe, and anterior cingulate cortices (Insausti & Muñoz, 2001; Cenquizca & Swanson, 2007; Risold et al., 1997; Van Groen & Wyss, 2003), as well as the rostrolateral NAc and rostral caudoputamen (Groenewegen et al., 1996; Naber & Witter, 1998; Swanson & Kohler, 1986). In contrast, non-human primate research indicates that the anterior hippocampus has stronger connectivity with the amygdala, hypothalamus, and medial (shell) of the NAc, VTA, Insula, and vmPFC (Poppenk et al., 2013). Critically, our findings could support a model in which different reward regions may engage discrete parts of the hippocampus to propagate downstream neural signals to distinct subcortical and cortical networks in service of adaptive behavior.

Our findings of distinct coupling amongst the hippocampus and reward regions set the foundation to begin investigating how individual differences in structural connectivity may differentially relate to individual differences in adaptive behavior. Another direction for future research is to investigate the functional significance of the heterogeneity in hippocampal connectivity profiles observed in this study by either examining brain-behavior relationships in large-scale studies or by conducting studies that simultaneously collect neuroimaging and DTI data. Finally, the probabilistic atlas generated in this study could be a valuable tool for guiding future research on the role of the hippocampus in psychopathology, particularly in conjunction with other neuroimaging techniques such as functional MRI and positron emission tomography (PET) in clinical populations. The hippocampus is disrupted in a variety of mental disorders, such as psychotic disorders (Lodge & Grace, 2011), posttraumatic stress disorder (Shin et al., 2006; van Rooij et al., 2015; Tanriverdi et al., 2022) and depression (Belujon & Grace, 2017; Grace, 2016), and is sensitive to environmental perturbations, such as childhood trauma and stress (Vythilingam et al., 2002; Kim & Diamond, 2002; Lupien & Lepage, 2001). In addition, childhood trauma and stress have been linked to anhedonia (lower reward functioning) in those at risk for psychosis (O’Brien et al., 2023), which highlights the clinical relevance to mapping out hippocampal contributions to reward circuitry.

The current study provides a foundation for future investigations into the anatomical and functional implications of the hippocampus and reward-related regions using more tailored regions of interest based on the underlying anatomical connectivity. This will enhance our understanding of the neural circuitry underlying adaptive behaviors and contribute to the development of novel therapeutic interventions for psychopathologies associated with reward processing (e.g. amotivation and anhedonia). The present study advances our knowledge of the structural connectivity of the HPC in humans, characterizing long-axis regions with distinct connectivity to reward-related regions.

